# A Nanoceutical Agent for Chemoprevention of Bilirubin Encephalopathy

**DOI:** 10.1101/2020.12.31.425023

**Authors:** Aniruddha Adhikari, Vinod K Bhutani, Susmita Mondal, Monojit Das, Soumendra Darbar, Ria Ghosh, Nabarun Polley, Anjan Kumar Das, Siddhartha Sankar Bhattacharya, Debasish Pal, Asim Kumar Mallick, Samir Kumar Pal

**Affiliations:** Department of Chemical, Biological and Macromolecular Sciences, S. N. Bose National Centre for Basic Sciences, Block JD, Sector 3, Salt Lake, Kolkata 700106, India; Department of Neonatal and Developmental Medicine, Lucile Packard Children’s Hospital, Stanford University, 750 Welch Road, Palo Alto, CA 94304, USA; Department of Zoology, Uluberia College, University of Calcutta, Uluberia, Howrah 711315, India; Department of Zoology, Vidyasagar University, Rangamati, Midnapore 721102, India; Research and Development Division, Dey’s Medical Stores (Mfg.) Pvt. Ltd., 62 Bondel Road, Ballygunge, Kolkata 700019, India; Technical Research Centre, S. N. Bose National Centre for Basic Sciences, Block JD, Sector 3, Salt Lake, Kolkata 700106, India; Physical Chemistry – innoFSPEC, University of Potsdam, Am Mühlenberg 3, 14476 Potsdam – Golm, Germany; Department of Pathology, Coochbehar Govt. Medical College and Hospital, Silver Jubilee Road, Coochbehar 736101, India; Department of Pediatric Medicine, Nil Ratan Sirkar Medical College and Hospital, 138 AJC Bose Road, Sealdah, Rajabazar, Kolkata 700014, India

**Author notes:** Corresponding author: **Prof. Samir Kumar Pal**, FNAE, Senior Professor, Department of Chemical, Biological and Macromolecular Sciences, S. N. Bose National Centre for Basic Sciences, Block JD, Sector 3, Salt Lake, Kolkata 700106, India.

**Keywords:** Acute bilirubin neurotoxicity, Severe Neonatal Hyperbilirubinemia, Chemoprevention, Nanotherapeutics, Nanotoxicity, Nanoceutical Pharmacokinetics

## Abstract

**Background:** Targeted degradation of bilirubin *in vivo* may enable safer and more effective approach to manage incipient bilirubin encephalopathy consequent to severe neonatal hyperbilirubinemia (SNH). This report builds on the use of a spinel structured mixed-valence transition metal oxide (trimanganese tetroxide) nanoparticle duly functionalized with biocompatible ligand citrate (C-Mn_3_O_4_ NP) having the ability to degrade bilirubin without photo-activation.

**Method:** The efficiency of C-Mn_3_O_4_ NP in *in vivo* degradation of serum bilirubin and amelioration of severe bilirubin encephalopathy and associated neurobehavioral changes was evaluated in C57BL/6j animal model of SNH.

**Results:** Oral single dose (0.25 mg kg^-1^ body weight) of the NPs efficiently reduced serum bilirubin levels (both conjugated and unconjugated) in study mice. It prevents bilirubin-induced neurotoxicity with reduction of SNH as observed by neurobehavioral and movement studies of SNH-mice. Pharmacokinetic data suggests intestinal reabsorption of the NPs and explain sustainable action. Biodistribution, pharmacokinetics, and biocompatibility of the NPs were tested during sub-chronic exposure.

**Conclusion:** Thus, we report preliminary studies that explore an affordable chemoprevention mechanism to acutely prevent or minimize bilirubin neurotoxicity in newborn infants.

**IMPACT STATEMENT:**

- Despite several attempts, no pharmaco-therapeutics are available for the treatment of severe neonatal hyperbilirubinemia (SNH) and associated neurotoxicity.
- Our newly developed nanodrug, citrate functionalized Mn_3_O_4_ nanoparticles (C-Mn_3_O_4_ NPs), can efficiently ameliorate SNH and associated neurotoxicity as investigated in preclinical rodent model.
- Chemoprevention effect of the nanodrug is found to be safe and sustainable.
- If successfully translated into clinical trials, C-Mn_3_O_4_ NPs could become the first drug to treat SNH.

## INTRODUCTION

Manganese (Mn), an essential trace mineral nutrient, is a cofactor for many crucial metabolic enzymes including manganese superoxide dismutase, arginase, and pyruvate carboxylase. Mn plays vital role in metabolism of amino acid, cholesterol, glucose, and carbohydrate; scavenging of reactive oxygen species (ROS); bone formation; reproduction; and immune response (1-4). Upon absorption, some of the Mn ions remain free, but remainder are mostly bound to transferrin, albumin, and plasma alpha-2-macroglobulin such that the circulating level ranges from 4-15 µg L^-1^ (5-8). These levels vary largely and do not serve as useful indicator of body Mn status (5). Overdose of Mn may lead to adverse neurological, reproductive, or respiratory effects, and has been reported among welders, miners, or persons with chronic inhalation exposure (9, 10). Despite the paradoxical behavior of this ‘Janus-faced’ metal in physiological systems, Mn oxides in nano-form (i.e., Mn_x_O_y_ nanomaterials) possess exciting redox-modulatory and catalytic properties necessary for novel therapeutic applications (1, 11-14).

Over the past decade, biomedical applications of nanometer-sized colloidal particles (i.e., nanoparticles) have been subject of intensive research due to their unique electronic, optical, and magnetic properties that are derived from the nanoscale dimensions and compositions (15). Numerous therapeutic and diagnostic modalities have also been developed using diverse array of nanomaterials (16). For example, Doxil® (doxorubicin liposomes), used for the treatment of AIDS-associated Kaposi’s sarcoma, was the first nanosized drug delivery system to receive United States Food and Drug Administration (US-FDA) approval (17). Similarly, Abraxane®, a 130 nm paclitaxel-decorated albumin was granted US-FDA approval as second-line treatment of breast cancer patients (18). Combidex® (ferumoxtran-10), an iron-oxide nanoparticle based imaging contrast agent is also being used in conjunction with magnetic resonance imaging (MRI) for differentiating cancer from normal lymph nodes (16). Recently, among numerous engineered nanomaterials manganese oxide nanomaterials showed promising results in treatment of chronic diseases like hepatic fibrosis (11), chronic kidney disease, neurodegenerative disorders (19) etc. in preclinical animal models. Our recent study demonstrated that manganese oxide nanoparticles do not display characteristic neurotoxicity (i.e., manganism), rather ameliorate Mn-induced neuralgic disorder (i.e., idiopathic Parkinson’s disorder) through mitochondrial protection and intracellular redox modulation (19). Interestingly in preliminary studies, the nanoparticle showed unprecedented catalytic activity towards bilirubin degradation without any photo- or chemo-activation in a controlled *in vitro* system (20).

The present study examines whether citrate functionalized trimanganese tetroxide nanoparticles (C-Mn_3_O_4_ NPs) can degrade bilirubin during *in vivo* administration and prevent bilirubin-induced neurotoxicity in a rodent model bred to exhibit severe neonatal hyperbilirubinemia (SNH).

## MATERIALS AND METHODS

### Rodent Model

All animal studies and experimental procedures were performed at Central Animal Facility, Uluberia College, India (Reg. No.: 2057/GO/ReRcBi/S/19/CPCSEA) following the protocol approved by the Institutional Animal Ethics Committee (Ref: 02/S/UC-IAEC/01/2019) as per standard guideline of Committee for the Purpose of Control and Supervision of Experiments on Animals (CPCSEA), Govt. of India. Non-diabetic C57BL/6j mice of both sex (age: 2-3 weeks, body weight, BW: 13±1.9 g) were used in the current study. Animals were housed under specific-pathogen-free (SPF) conditions (maximum 5 mice per polypropylene cage; temp: 20±1.5° C; ∼55% relative humidity) under a 12-h light and 12-h dark cycle with access to food (standard chow for mice, Saha Enterprize, India) and water *ad libitum*. Autoclaved nest material and paper houses served as cage enrichment for mice. Animal cages were always randomly assigned to treatment or control groups.

### *In vivo* Chemoprevention

The study was conducted in two phases. In the first phase, the *in vivo* bilirubin degradation ability of C-Mn_3_O_4_ NPs was tested in time dependent manner along with plasma pharmacokinetics (PK), and pharmacodynamics (PD). In the second phase, sustainability of the treatment and potential chemoprevention effect against bilirubin induced neurotoxicity were evaluated. For induction of SNH, the well-known phenyl hydrazine (PHz) intoxication model was used (21, 22). PHz drastically breaks down hemoglobin inside the rodent body, in turn, simulates a pathological condition of SNH and hemolytic anemia. One of the major reasons for choosing this particular model is the non-involvement of liver in the whole pathophysiology.

#### Phase 1

Mice were randomly divided into five groups (N=10/group). Animals of Group 1 served as control and received normal saline (150 µL; oral). Animals of Group 2 – 4 received two doses (i.p.) of PHz; 60 mg kg^-1^ BW at day-1 and 30 mg kg^-1^ BW at day-3. Animals of Group 3 were further treated with C-Mn_3_O_4_ NPs (0.25 mg kg^-1^ BW; oral) at day-3. Group 4 animals received citrate (0.15 mg kg^-1^ BW; oral) at day-3. Animals of Group 5 served as NP control and administered with C-Mn_3_O_4_ NPs (0.25 mg kg^-1^ BW; oral) at day-3. In phase-1, all measurements were performed on day-3.

#### Phase 2

Animals were randomly divided into five groups. Animals of Group 1 (N=10) served as control and received normal saline (150 µL; oral). Animals of Group 2 – 4 (N=30/group) received three doses (i.p.) of PHz; 45 mg kg^-1^ BW at day-1 and 30 mg kg^-1^ BW at day-3 and day-5 for induction of SNH. At day-1, −3, and −5, animals of Group 3 were further treated with C-Mn_3_O_4_ NPs (0.25 mg kg^-1^ BW; oral). Group 4 animals received citrate (0.15 mg kg^-1^ BW; oral) at day-1, −3, and −5. Group 5 animals were administered with C-Mn_3_O_4_ NPs (0.25 mg kg^-1^ BW; oral) at day-1, −3, and −5. In phase-2, all biochemical and neurobehavioral measurements were performed on day-6.

The time gap between two treatments on same day was ∼1 h. PHz was always administered first. The optimum doses (both for intoxication and therapy) as well as treatment protocol was designed based on published literature and prior pilot studies performed by us.

### Bilirubin Estimation from Serum Samples

For biochemical studies, blood samples were collected from retro-orbital plexus in non-heparinized sterile tubes just before sacrifice. After collection of blood, serum was instantly separated by centrifugation (3500 rpm; 10 mins) at 4° C, and in dark condition. Bilirubin measurements were completed within 30 mins of serum isolation. Care was taken to keep the samples at 4° C away from light exposure before beginning of measurements. Total serum bilirubin (TSB), and unconjugated bilirubin (UCB) were measured following di-azo method using commercially available test kits (Autospan Liquid Gold, Span Diagnostic Ltd., India), and double beam UV-Vis absorbance spectrometer (Model UV-Vis 2600, Shimadzu, Japan). Precision of the instrument was 0.001 OD. Each time instrument was calibrated using standard bilirubin solution (5.0 mg dL^-1^). Variation of the measurements lied within 2 – 3%.

### Histopathology

For histopathological investigation, brains were removed from the sacrificed animals, and fixed in 10% buffered formalin for upto 7 days. Paraffin embedded sections (5-8 µm) were stained with hematoxylin and eosin following standard method.

### Behavioral Studies

#### Open field test (OFT)

At day-6 of phase-2, animals were first acclimatized to the dimly lit experimental room (∼15 lx) for 30 min and then individually placed in an illuminated open field apparatus (45 cm × 45 cm × 45 cm; ∼1200 lx). They were permitted to explore freely for 5 min after an initial 1-min habituation phase. Movement pattern and associated parameters was analyzed from videos recorded during experiments.

#### Gait analysis

For gait measurement, the forelimbs and hindlimbs of each mouse were coated with red and blue nontoxic paints, respectively. Then they were individually placed on the walkway (30 cm × 8 cm; closed sidewise), and allowed to move freely in both directions. Before final experiments, animals were allowed to habituate and cross the walkway 2 times. The gait parameters were calculated from the footprints imprinted on a sheet of white paper which was placed on the floor of the walkway.

#### Balance beam test

The ability of the animals to pass through a narrow wooden beam (1 cm × 80 cm; elevated 1 m above the floor) to reach a dark box was evaluated in balance beam test. A white light (∼1200 lux) illuminated the origin to force the mice to cross the beam. The time required to reach the target box, and the number of forelimb and hind-limb paw slips were recorded. A paw slip was defined as any paw coming off the top of the beam or any limb use on the side of the beam. Final data were recorded after 3 habituation trial.

#### Pole test

Individual animals were placed facing upward on the top of a vertically standing wooden pole (1 cm × 80 cm) whose base was positioned in the mouse home cage and titled at 45° to stand on a nearby wall. Mice were placed with their heads directed upwards on the upmost part and were forced to attempt descending the pole to enter the home cage. The time required for mice to descend and reach the floor with all four paws were recorded.

### Pharmacokinetics

For PK studies, animals were administered with C-Mn_3_O_4_ NPs (0.25 mg kg^-1^ BW; oral). Then, blood was collected at different time points, and Mn contents were estimated using inductively coupled plasma atomic emission spectroscopy (ICP-AES) (ARCOS-Simultaneous ICP Spectrometer, SPECTRO Analytical Instruments GmBH, Germany). Open acid digestion method was employed for preparation of samples. In brief, dried tissue samples (in liquid nitrogen) were dissolved in 3:2:1 mixture of HNO_3_ (Merck, Germany), H_2_SO_4_ (Merck, Germany), and H_2_O_2_ (Merck, Germany), and heated at 150° C until only a residue remained. Then, the residues were diluted to 10 mL using Mili-Q water.

### Statistical Analysis

Kaplan-Meier survival curves were used to illustrate mortality due to SNH after PHz intoxication, and NP co-treatment. Differences in survival between groups were assessed by the log-rank test with multiple pair-wise comparisons performed using the Mantel-Cox method. All quantitative data are expressed as Mean ± Standard Deviation (SD), unless otherwise stated. Unpaired *t* test with Welch’s correction was used to compare between two groups. One-way analysis of variance (ANOVA) followed by Tukey’s *post hoc* multiple comparison test was performed for comparison between multiple groups. Beforehand, the normality of each parameter was checked by normal quantile–quantile plots. Sample size in our animal studies were determined following the standard sample sizes previously been used in similar experiments as per relevant literature. For some experiments, sample size was adapted to the observed effect size, and numbers were increased to 15–20 animals per group. Designated sample size (in figure legends) always refers to biological replicates (independent animals). GraphPad Prism v8.0 (GraphPad Software), and Sigmaplot v14.0 (Systat Software, Inc.) were used for statistical analysis. For all comparisons, a *P* value <0.05 was considered statistically significant.

## RESULTS

Effect of C-Mn_3_O_4_ NPs on unconjugated bilirubin (UCB) level was monitored at 2 h interval upto 24 h, after treatment with single oral dose (0.25 mg kg^-1^ BW) of C-Mn_3_O_4_ NPs, once UCB level reached 4.78±0.34 mg dL^-1^ (compared to 0.22±0.05 mg dL^-1^ of control; *P* <.0001, two-tailed *t*-test), and TSB level reached 6.13±0.44 mg dL^-1^ (compared to 0.29±0.05 mg dL^-1^ of control; *P* <.0001, two-tailed *t*-test) due to PHz intoxication. As the results suggest, C-Mn_3_O_4_ NPs was able to irreversibly decrease both UCB (0.62±0.18 mg dL^-1^; *P* <.0001, two-tailed *t*-test) (Figure 1a) and TSB levels (1.15±0.22 mg dL^-1^; *P* <.0001, two-tailed *t*-test) (Figure 1b) to normal range within 12 h. Fitting the UCB degradation kinetic data in logistic regression model (adj. *R*^*2*^ 0.994), the time required for 50%, and 80% bilirubin degradation were found to be 6.6 h, and 8.3 h, respectively (Figure 1a). Similarly, for TSB the time required for 50%, and 80% bilirubin degradation were 7.3 h, and 10.6 h, respectively (adj. *R*^*2*^ 0.996) (Figure 1b). It is worth mentioning here that after oral exposure ∼4 h is required for the NPs to reach sufficient plasma concentration (see PK-PD section for details). Thus, effectively within 3 h of absorption, the C-Mn_3_O_4_ NPs were able to degrade 50% of the UCB, and ∼45% of TSB *in vivo*. While comparing the time dependent changes in UCB, TSB, and direct bilirubin (DB) levels, we found lesser efficiency of the C-Mn_3_O_4_ NPs towards degradation of DB (∼70%; *P* <.0001, one-way ANOVA) compared to UCB (∼96%), and TSB (∼92%) at 24 h (Supplementary Figure S1). Treatment with citrate (0.15 mg kg^-1^ BW; oral) after PHz-intoxication did not affect the UCB (6.27±1.66 mg dL^-1^; *P*=.9071 compared to PHz-treated, one-way ANOVA, F (4, 45)=12.57), and TSB levels (7.19±1.53 mg dL^-1^; *P*=.9944 compared to PHz-treated, one-way ANOVA, F (4, 45)=10.53) (Figure 1a-inset & 1b-inset). Cumulatively, these results indicate that C-Mn_3_O_4_ NPs can efficiently degrade bilirubin *in vivo* and, the bilirubin degradation observed after C-Mn_3_O_4_ NP treatment is due to the catalytic nature of the nanoceutical complex itself, not because of the outer citrate coating.

**Figure 1.**
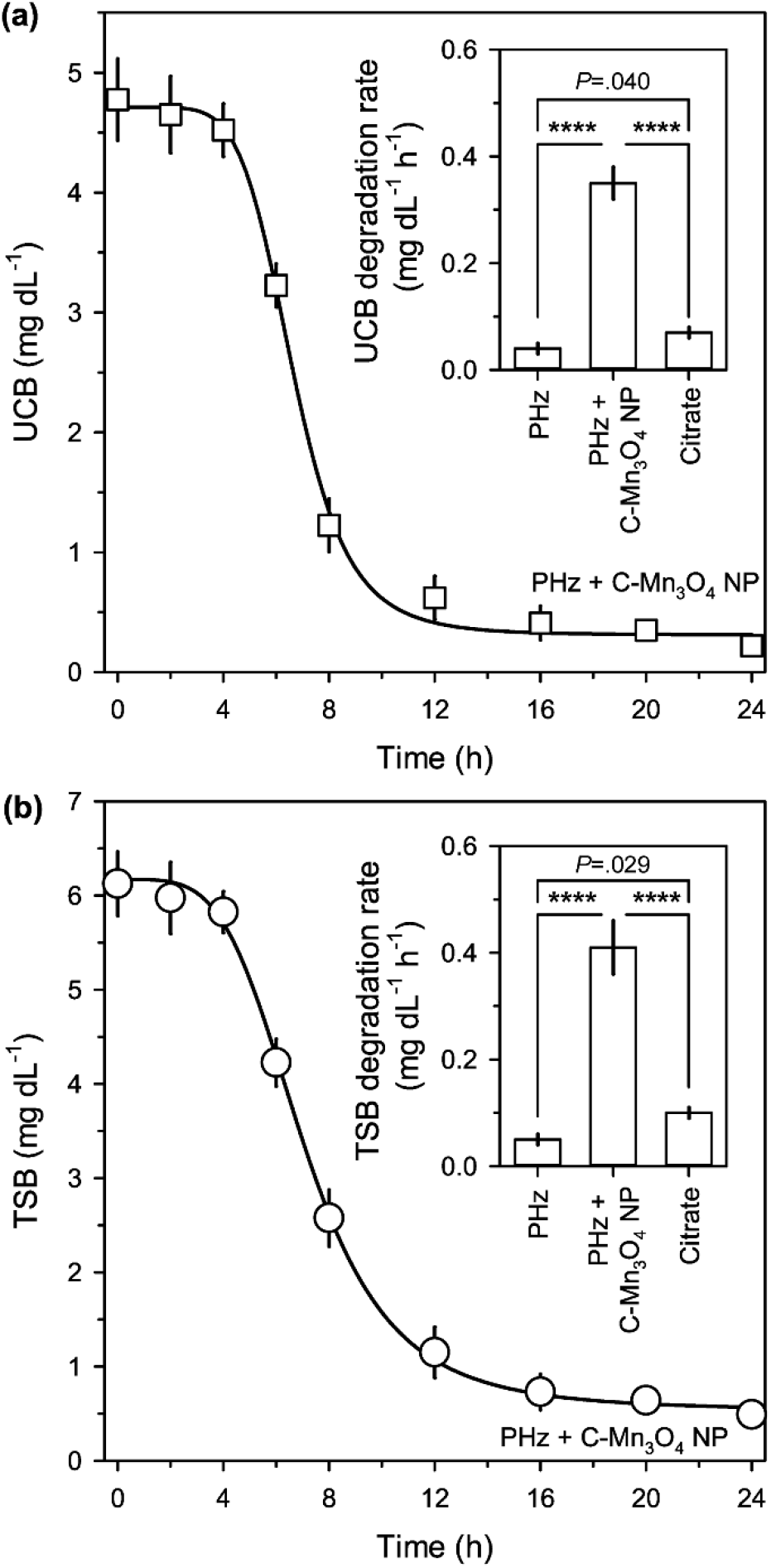
*In vivo* bilirubin degradation kinetics after treatment with C-Mn_3_O_4_ NPs in PHz-induced hyperbilirubinemia mice. (a) Unconjugated bilirubin (UCB). (b) Total serum bilirubin (TSB). Insets show bilirubin degradation rates across different treatment groups. Error bars represent standard deviation from the mean (*n* = 6). **** signifies *P*<0.0001, one-way ANOVA, Tukey’s Multiple Comparison Test (*post-hoc*).

Duration of the chemoprevention effect was evident from our phase-2 study. PHz drastically increased both UCB (6.69±1.56 mg dL^-1^; *P* <.0001, F (2,15)=45.96, one-way ANOVA), and TSB (7.39±1.65 mg dL^-1^; *P* <.0001, F (2,15)=58.00, one-way ANOVA) compared to control (UCB: 0.22±0.05 mg dL^-1^; TSB: 0.29±0.05 mg dL^-1^) (Figure 2a & 2b) simulating a condition of SNH. Reduction in hematocrit (Hct) value further confirmed induction of PHz-induced hemolytic condition (Supplementary Figure S2a). Co-treatment with C-Mn_3_O_4_ NPs (0.25 mg kg^-1^ BW; oral) significantly reduced both UCB (0.31±0.11 mg dL^-1^), and TSB (0.42±0.14 mg dL^-1^) compared to PHz-induced untreated littermates (*P* <.0001; one-way ANOVA) (Figure 2a & 2b). We further monitored the UCB and TSB levels for additional one week once the treatment ended (Supplementary Figure S2b-c). There was no recurrent increase in UCB or TSB during the one week period, indicating sustainability of the chemoprevention by C-Mn_3_O_4_ NP. Animals co-treated with citrate (0.15 mg kg^-1^ BW; oral) + PHz showed UCB (6.27±1.66 mg dL^-1^) and TSB (7.19±1.53 mg dL^-1^) levels similar to that of PHz-intoxicated group (UCB: 6.69±1.55 mg dL^-1^, *P*=.8457; TSB: 7.39±1.65 mg dL^-1^, *P*=.9653, one-way ANOVA) (Figure 2a & 2b). The successful reduction of UCB and TSB by oral administration further suggest that C-Mn_3_O_4_ NPs with high stability and gastric passage will be of potential easy use in human neonates too.

**Figure 2.**
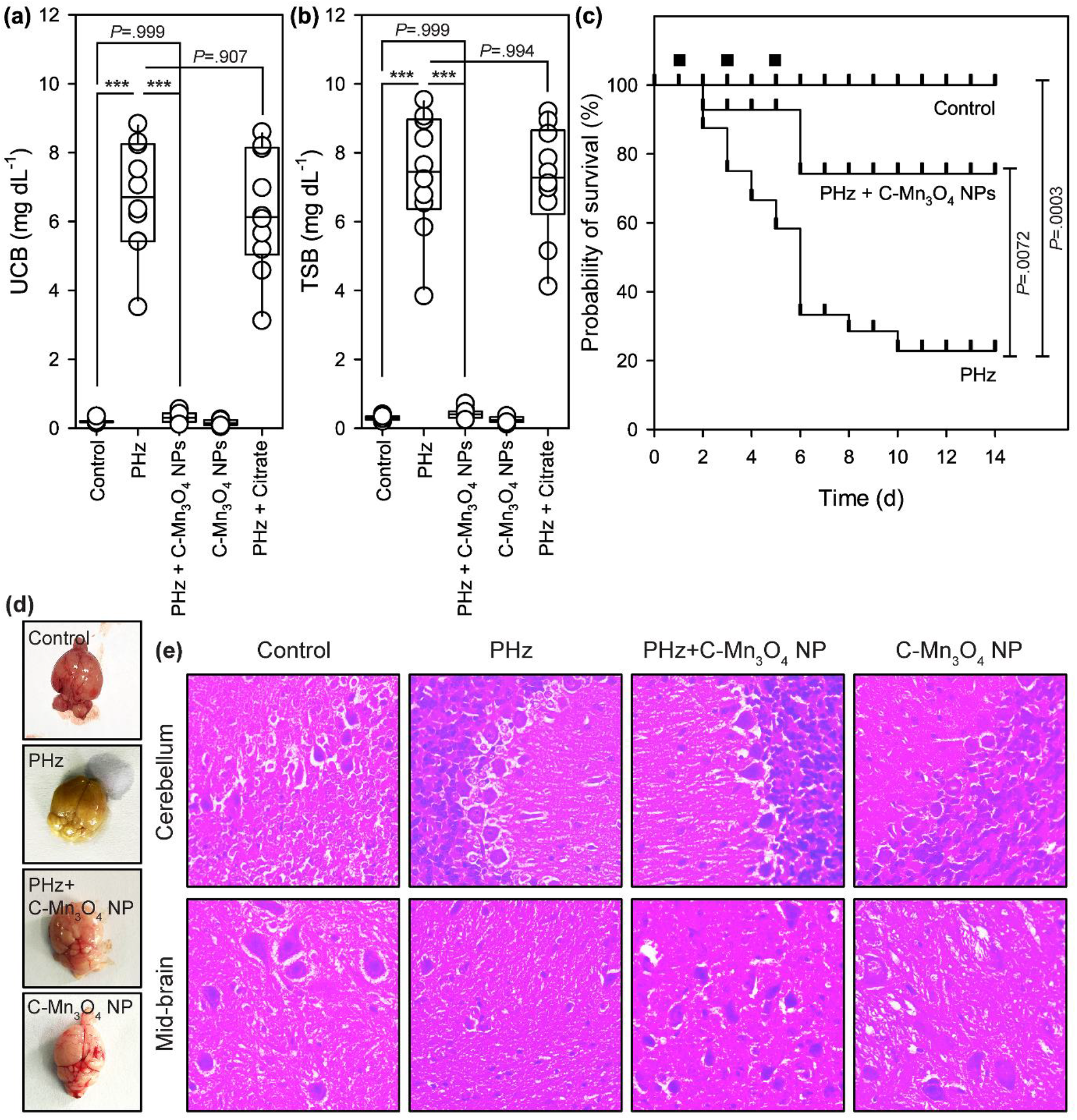
Effect of C-Mn_3_O_4_ NPs on severe neonatal hyperbilirubinemia and associated neurotoxicity. (a) Unconjugated bilirubin level. (b) Total serum bilirubin level. (c) Probability of survival of differently treated mice as a function of time (Kaplan–Meier analysis). The black squares represents days of treatment. (d) Photograph of isolated brains. (e) Micrographs of hematoxylin and eosin stained brain sections. *** signifies *P*<0.001, one-way ANOVA, Tukey’s Multiple Comparison Test (*post-hoc*). *n*=6.

Repeated dose PHz administration induced significant lethality in the study mice (Hazards Ratio, HR (Mantel-Haenszel): 0.158; 95% CI of HR: 0.058-0.32; log rank χ^2^ (Mantel-Cox): 12.94; df: 1; *P*=.0003) (Figure 2c). Nearly 50% of the animals died during the first week due to PHz-induced SNH (median survival: 6 days). Strikingly, C-Mn_3_O_4_ NP treatment was not only well tolerated by the study mice (Group 5; No mortality) but also significantly reduced lethality in PHz-intoxicated C-Mn_3_O_4_ NP co-treated mice (Group 3; 10% mortality) as compared to the untreated PHz-intoxicated group (Group 2; 60% mortality) (HR (Mantel-Haenszel): 3.650; 95% CI of HR: 1.419-9.391; log rank χ^2^ (Mantel-Cox): 7.210; df: 1; *P*=0.0072). The difference in the survival rates of PHz + C-Mn_3_O_4_ NP co-treated mice and PHz-intoxicated untreated mice was most notable during the first week of the treatment (Figure 2c). Necropsy revealed, consistent with the blood biochemical analysis, bilirubin induced acute neurotoxicity and extreme hemolysis as the cause of death. Therefore, our study evidently suggest that C-Mn_3_O_4_ NPs can reduce the severity and mortality of SNH through reduction of bilirubin level in experimental rodent model.

To induce a kernicterus-like syndrome, prolonged repeated PHz administration was used, and *in vivo* bilirubin-induced neurotoxicity in the PHz-treated animals are evident from the digital photograph of isolated brains (Figure 2d). The intense yellow discoloration, attributed to bilirubin, in gross brain photograph of PHz-induced rodents indicated accumulation of bilirubin in the brain. This yellow discoloration was not observed in C-Mn_3_O_4_ NP + PHz co-treated group, indicating low accumulation of neurotoxic bilirubin in brain. In accordance with the biochemical results, histopathological investigations of the isolated brain sections revealed severe bilirubin-induced damage to the PHz-intoxicated animals (Figure 2e and Supplementary Figure S3). In cerebellum, marked reduction (∼35%) in number of cells per field, a hallmark of bilirubin induced severe neurotoxicity was noticed in PHz-intoxicated animals. Number of Purkinje neurons (∼30%) and cells of substantia nigra (∼40%) were also decreased significantly compared to control. Other pathological features include extensive fibrosis, spongiosis (vacuolation) in brain parenchyma (linked to depletion of neurons), shrinkage in cell size, eosinophilic neurons (characteristic of degenerating cells) and gliosis. In contrast, the PHz+C-Mn_3_O_4_ NP co-treated animals showed normal cellular architecture with mild gliosis, remnant of PHz-induced brain damage. Number of cells per field in both cerebellum and mid-brain region were comparable to control animals. No shrinkage of cells were observed. C-Mn_3_O_4_ NP treated animal showed brain architecture similar to those of untreated control. In summary, the histopathological examinations illustrate the protective action of C-Mn_3_O_4_ NPs at cellular level to ameliorate bilirubin-induced neurotoxicity.

In view of chronic sequelae, i.e., motor dysfunction and ataxia, we tested the performance of the animals in OFT (Figure 3a). The measured total distance moved (Figure 3b) by affected rodents, and the time spent at the center of the field (Figure 3c) was less in PHz-treated group compared to control (Total distance moved: 41.7±5.1 m vs. 47.8±3.4 m, *P*=.0112, F (2, 15)=13.24, one-way ANOVA; Time spent at the center: 16.2±4.1% vs. 29.1±4.7%; *P*=.0007, F (2, 15)=6.99, one-way ANOVA) and PHz+C-Mn_3_O_4_ NP co-treated group (Total distance moved: 41.7±5.1 m vs. 48.2±4.4 m, *P*=.0068; Time spent at the center: 16.2±4.1% vs. 27.2±5.2%; *P*=.0028, one-way ANOVA), and illustrate serious movement disorders. In addition, the time to cross a beam (Figure 3d: *P*=.0018, F (2, 15)=9.29, one-way ANOVA), and time to descend a pole (Figure 3e: Time to descend: *P*<.0001, F (2, 15)=34.93, one-way ANOVA) were also higher in PHz-induced group to indicate severe motor dysfunction, one of the characteristic features of bilirubin-induced severe neurotoxicity. Treatment with C-Mn_3_O_4_ NP was protective against motor dysfunction injury (Time to cross: *P*=.0532; Time to descend: *P*<.0001, one-way ANOVA). To get further insight, we monitored movement patterns of the animals (Figure 3f). The results show a significant decrease in stride length (Figure 3g: 3.83±0.15 cm vs. 2.78±0.14 cm; *P*<.0001, two-tailed *t*-test) and stride width (Figure 3h: 2.89±0.11 cm vs. 1.92±0.13 cm; *P*<.0001, two-tailed *t*-test) in PHz-induced group. Stance length (Figure 3i: 3.19±0.12 cm vs. 2.55±0.14 cm; *P*<.0001, two-tailed *t*-test), and stance angle (Figure 3j: 44.2±2.1° vs. 20.6±1.2°; *P*<.0001, two-tailed *t*-test) were also significantly decreased in the PHz-treated group. Use of C-Mn_3_O_4_ NPs allowed the rodents to retain their regular movement pattern (Figure 3f-3j; *P*<.0001 compared to PHz-intoxicated group for all parameters, two-tailed *t*-test), similar to the control animals.

**Figure 3.**
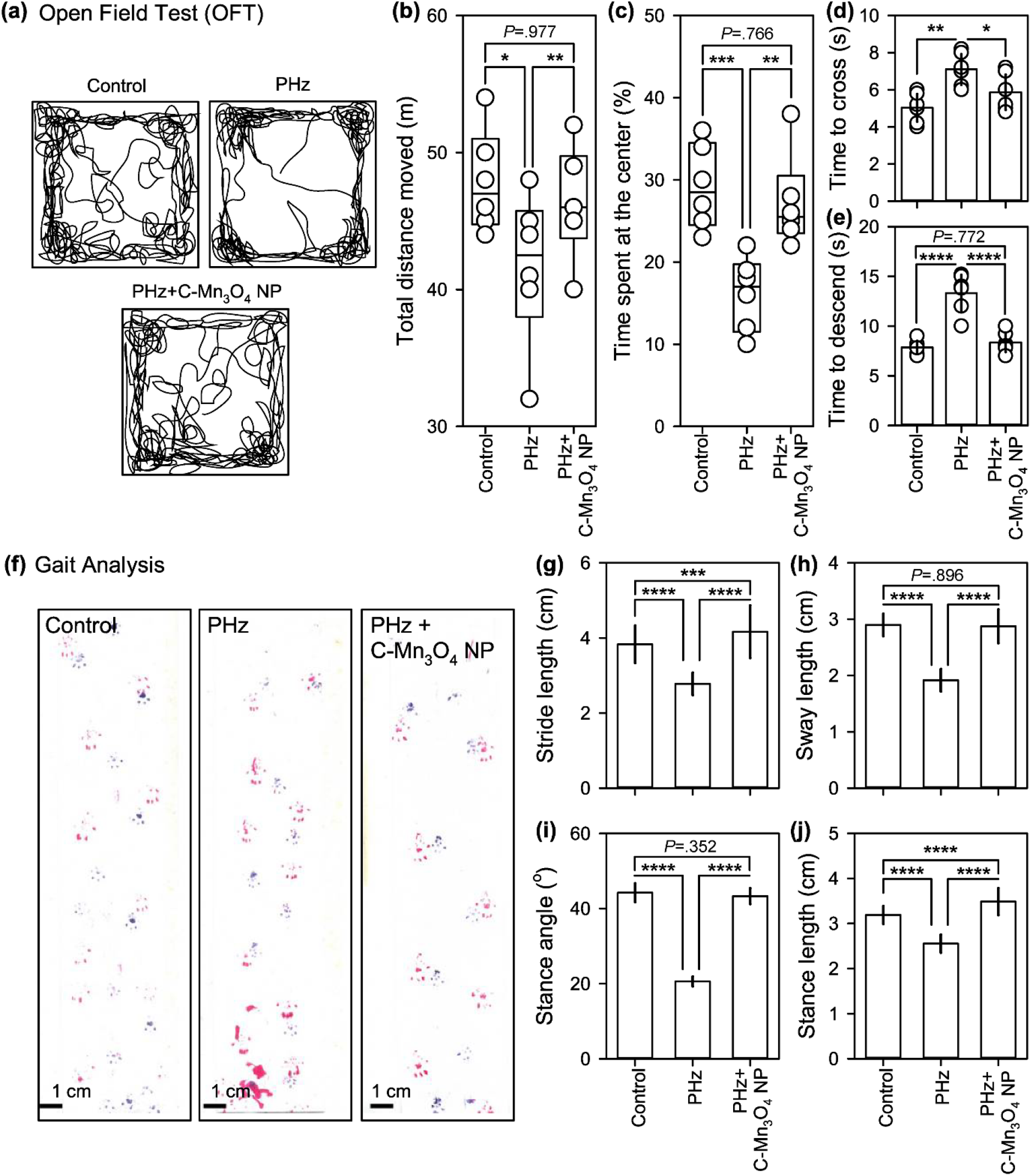
Effect of C-Mn_3_O_4_ NPs on severe neonatal hyperbilirubinemia induced neurobehavioral changes. (a) Trace of open field activity. (b) Total distance moved in open field apparatus. (c) Time spent at the center of the open field. (d) Time to cross a beam. (e) Time to descend a pole. (f) Trace of movement. (g) Stride length. (h) Sway length. (i) Stance angle. (j) Stance length. Error bars (except box and whisker plots) represent standard deviation from the mean (*n* = 6). *, **, ***, **** signifies *P*<0.05, *P*<0.01, *P*<0.001, and *P*<0.0001 respectively, one-way ANOVA, Tukey’s Multiple Comparison Test (*post-hoc*).

Pharmacokinetics and pharmacodynamics dictated by translocation and biodistribution of C-Mn_3_O_4_ NPs are the two major factors that mediate *in vivo* therapeutic activity. Time dependent plasma concentration profile of C-Mn_3_O_4_ NPs indicating its absorption and elimination is reported after oral delivery of a 0.25 mg kg^-1^ BW dose of C-Mn_3_O_4_ NP in plasma Mn concentration versus time plot (Supplementary Figure S4a). The PK parameters, calculated by a non-compartmental approach, yield a maximum plasma concentration (C_MAX_) of 1.71±0.25 µg mL^-1^ of Mn for the C-Mn_3_O_4_ NPs at (t_MAX_) 12.1±0.2 h (Supplementary Table S1). The plasma area under the curve (AUC) was 18.01±1.47 µg mL^-1^ h with a clearance of 12.3±0.8 L h^-1^ kg^-1^. A steady low concentration of NPs (Mn ∼0.1 µg mL^-1^) was maintained in plasma throughout 48 h window of experiment. The mean plasma concentration curve presented three-peak absorption phases (at ∼1.0 h, ∼6.0 h, and 12.0 h) indicated as I, II, and III in Supplementary Figure S4a. The deconvoluted spectra shows a significant overlap between intestinal absorption and hepato-biliary reabsorption, which resulted in the higher third peak (∼12 h). The first peak may be due to the absorption in the upper gastro-intestinal tract, as direct delivery of C-Mn_3_O_4_ NPs to the stomach by oral gavage resulted in disappearance of the first peak (Supplementary Figure S4b). Urinary Mn-levels were below detectable range upto 48 h. Mn content of feces are illustrated in Supplementary Figure S4c with maximum concentration (C_MAX_ = 0.68±0.07 µg g^-1^) at ∼18.0 h.

The effect of C-Mn_3_O_4_ NPs on bilirubin degradation are correlated with plasma C-Mn_3_O_4_ NP concentration profile (Supplementary Figure S4d). The results show an inverse relationship between UCB and plasma C-Mn_3_O_4_ NP concentration. UCB levels plotted against plasma C-Mn_3_O_4_ NP concentration (Supplementary Figure S4e & 4f) describes a dose dependent correlation (adj. *R*^*2*^ 0.860; non-linear logistic regression model) suggestive of *in vivo* catalytic degradation of bilirubin by C-Mn_3_O_4_ NPs.

## DISCUSSION

Our study demonstrates that the nanoceutical compound C-Mn_3_O_4_ NPs can degrade bilirubin during *in vivo* administration and prevent bilirubin induced neurotoxicity in a rodent model of SNH. A single oral dose reduces total as well as unconjugated bilirubin level and importantly, the nanoceutical agent has the capacity to significantly decrease or reverse early bilirubin neurotoxicity. These signs are described by impaired motor functions as observed in OFT, motor movements and gait analysis in PHz intoxicated group. The movement was less in OFT, and significant decrease was observed in step length, and step width in gait analysis. The motor function as measured using beam traversal, and pole test were also less. In addition, stance length, and stance angle were significantly decreased in the study group. We report that administration of C-Mn_3_O_4_ NPs facilitated the rodents to retain their normal movement pattern and sensory motor function. The histopathological examination reveals that C-Mn_3_O_4_ NPs can protect the neural cells (particularly of mid-brain and cerebellum region of the brain) from bilirubin-induced severe damages. The pharmacokinetics and pharmacodynamics studies further indicates the sustainable therapeutic action of this novel agent. The intestinal reabsorption after hepatobiliary recirculation was found to be important in maintaining longer blood circulation time of C-Mn_3_O_4_ NPs. Within 24 hours the materials is completely excreted from blood, the major route of excretion was found to be through feces.

Mn_3_O_4_ nanoparticles are mixed-valence transition metal oxides with a spinel structure, and are an important class of nanomaterials that have been investigated extensively (11-14, 19, 20). They primarily serve as catalysts due to their efficient activity, low cost, simple preparation, and high stability (23). They have become one of the key topics in contemporary research because of their excellent bifunctional oxygen electrode activity, photocatalytic efficiency and potential applications in several redox reactions (20). Manganese oxide-based NPs have emerged as potent MRI contrast agents owing to their impressive contrast ability, low toxicity, and longer circulation time in the bloodstream compared to the conventional gadolinium (Gd) or iron oxide based probes (24, 25). Successful *in vivo* use of manganese oxide nanomaterials as diagnostic agents paved the way to their therapeutic use. In 2016, our group for the first time has demonstrated that citrate functionalized trimanganese tetroxide nanoparticles (C-Mn_3_O_4_ NPs) can be administered orally to treat hepatic fibrosis in a rodent model (11). It is worth mentioning that chronic liver diseases (including fibrosis) represent a major global health problem both for their high prevalence worldwide and, in the more advanced stages, for the limited available curative treatment options (26, 27). In fact, when lesions of different etiologies chronically affect the liver, triggering the fibrogenesis mechanisms, damage has already occurred, and the progression of fibrosis will have a major clinical impact entailing severe complications, expensive treatments and death in end-stage liver disease (28). Recently, we have showed that the C-Mn_3_O_4_ NPs do not possess the characteristic manganese mediated neurotoxicity, rather it ameliorates Mn-induced Parkinson’s like syndrome (19). Therefore, it is safe for *in vivo* administration. Moreover, we have studied the systemic toxicity of C-Mn_3_O_4_ NPs upon 90 days of repeated dose chronic exposure in C57BL/6j mice. The preliminary results revealed no signs of toxicity at therapeutic dose, further supporting the biocompatibility of the nanomedicine.

Previously in controlled *in vitro* laboratory settings, we have demonstrated that C-Mn_3_O_4_ NPs has unprecedented catalytic activity towards degradation of bilirubin without any photo-activation (20). The *in vivo* bilirubin degradation ability of C-Mn_3_O_4_ NPs in SNH rodent model can be attributed to its inherent redox behavior. Briefly, in binary spinel Mn_3_O_4_ NPs all the tetrahedral A sites hold a divalent cation, Mn^2+^ (3d^5^) whereas all the octahedral B sites are occupied by trivalent cations, Mn^3+^ (3d^4^) (29). Our preliminary *in vitro* studies showed that the spontaneous comproportionation and disproportionation of surface Mn-ions lead to exceptional redox activity of the NPs (20, 30). The redox activity together with the ligand to metal charge transfer (LMCT) from the coordinating ligand citrate to NP surface results into the unprecedented bilirubin degradation ability (12, 20). The degraded product was identified as methyl-veny-maleimide (MVM), one of the well-known oxidative breakdown products of bilirubin (31). These initial findings alluded us to explore the potential clinical use of C-Mn_3_O_4_ NPs for human subjects at risk for bilirubin neurotoxicity, which have led us to consider a novel *in vivo* approach in a rodent model of neonatal hyperbilirubinemia. This is particularly important from pediatric point of view to find a drug that degrades bilirubin *in vivo* and can alter the subsequent trajectory of bilirubin rate of rise has the potential to protect infants from severe neonatal hyperbilirubinemia because each year, at least 0.5 million term or near-term newborn infants are affected with severe hyperbilirubinemia (TSB>25 mg/dl), of whom 0.15 million die and over 0.07 million sustain moderate or severe disability (32).

These preliminary studies have to be taken in context with other metallic chemopreventive agents, notably metalloporphyrins (such as, tin and zinc mesoporhyrin) reduces bilirubin production and bilirubin levels in human newborns through competitive inhibition of heme oxygenase (HO), the rate-limiting enzyme in the catabolism of heme to bilirubin, and therefore inhibits formation of bilirubin (33, 34, 35). This approach compares to facilitated bilirubin elimination by phototherapy or an exchange transfusion. After 3 decades of extensive and comprehensive scholarly and industry sponsored studies, the FDA studied the submission of clinical translation data and has declined the industry’s request for license.

A limitation of the study is the use of chemically induced rodent model of SNH. Although, PHz-induced mice model is one of the standard and most widely used animal model for testing efficacy of drugs against SNH (21, 22), evaluating the efficacy of C-Mn_3_O_4_ NPs in genetically modified animals like Gunn rat can provide more insight into the therapeutic mechanism. Administration of C-Mn_3_O_4_ NPs in combination with phototherapy could be more effective, and needs to be studied further. However, to the best of our knowledge our robust preclinical study provides strong support to the hypothesis that C-Mn_3_O_4_ NPs could effectively reduce SNH when administered orally and possibly reverse acute bilirubin neurotoxicity.

## CONCLUSION

We report the first, to our knowledge, a novel chemoprevention nanoceutical agent that selectively degrades bilirubin *in vivo* and likely to ameliorate acute bilirubin neurotoxicity. Our data primarily provides direct evidence that single oral administration of citrate functionalized Mn_3_O_4_ nanoparticles (C-Mn_3_O_4_ NPs) can reduce severe neonatal hyperbilirubinemia (SNH) in a rodent model. We documented an alternate catalytic-therapeutic mechanism that governs targeted disruption of bilirubin metabolism. Robust properties of nanoparticles were studied to demonstrate stability in acidic conditions, thus protected from low pH of stomach that would enable lysosomal degradation and oral administration. This novel nano-particle, C-Mn_3_O_4_ NPs may have the potential to become an affordable therapeutic option for the newborn infant who may be at the risk for acute bilirubin neurotoxicity. Risk-benefit studies are warranted prior to any clinical application or inquiry.

## ACKNOWLEDGEMENT

MD thanks University Grants Commission (UGC), Govt. of India for Junior Research Fellowship. SKP thanks the Indian National Academy of Engineering (INAE) for the Abdul Kalam Technology Innovation National Fellowship, INAE/121/AKF. The authors thank the Department of Biotechnology (DBT, West Bengal) for the financial grant under BOOST scheme, 339/WBBDC/1P-2/2013.

## DATA AVAILABILITY

All data that support the findings of this study are available within the published article (and supplementary information files). Study related additional data will be available from SKP upon legitimate request.

## AUTHOR CONTRIBUTION

A.A., V.K.B., and S.K.P. conceived and planned the experiments. A.A., S.M., M.D., N.P., and S.D. performed animal experiments and biochemical studies. A.A. executed numerical calculations and statistical analysis. A.A., S.D., S.S.B., D.P., R.J.W., A.K.M., V.K.B and S.K.P. contributed to interpretation of results. A.K.D. carried out the histological studies and analyzed the data. A.A. and V.K.B. took the lead in writing the manuscript. All authors provided critical feedback and helped in shaping the research, analysis, interpretation and writing. All authors reviewed, edited and approved the final version of the manuscript.

## CONSENT STATEMENT

This study does not involve human subject or clinical study. Thus, patient consent was not required. All animal studies and experimental procedures were performed at Central Animal Facility, Uluberia College, India (Reg. No.: 2057/GO/ReRcBi/S/19/CPCSEA) following the protocol approved by the Institutional Animal Ethics Committee (Ref: 02/S/UC-IAEC/01/2019) as per standard guideline of Committee for the Purpose of Control and Supervision of Experiments on Animals (CPCSEA), Govt. of India.

