## Supplementary Information for "A Nanoceutical Agent for Chemoprevention of Bilirubin Encephalopathy"

*Corresponding author:

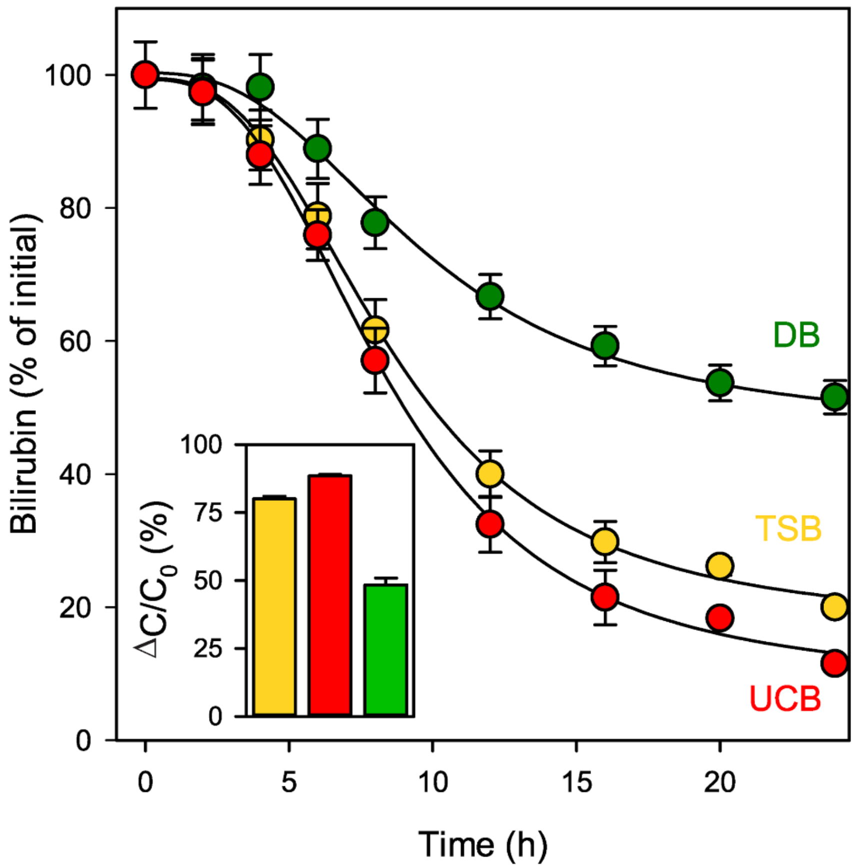


**Supplementary Figure S1. Comparison of *in vivo* bilirubin degradation kinetics for different types of bilirubin in PHz+C-$Mn_3O_4$ NP co-treated group.** Data represented as percentage (%) of initial concentration. Inset shows % degradation of different bilirubins (UCB, TSB, and DB) after 12 h of treatment. Error bars represents standard deviation (SD) from the mean (*n* = 6).

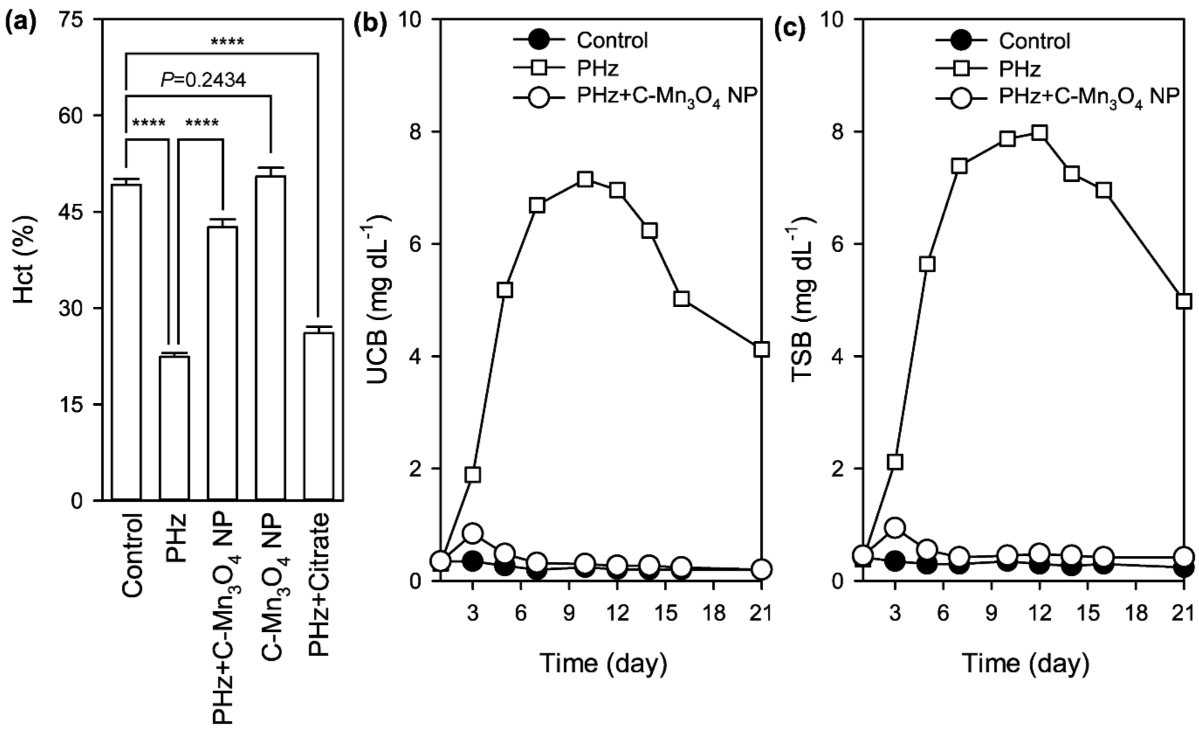


**Supplementary Figure S2.** (a) Hct values across treatment groups. (b) & (c) Sustained action of C-$Mn_3O_4$ NPs for amelioration of severe neonatal hyperbilirubinemia. Data represented as Mean±SD ($n = 6$). *, **, ***, **** signifies $P<0.05$, $P<0.01$, $P<0.001$, and $P<0.0001$ respectively, one-way ANOVA, Tukey's Multiple Comparison Test (*post-hoc*).

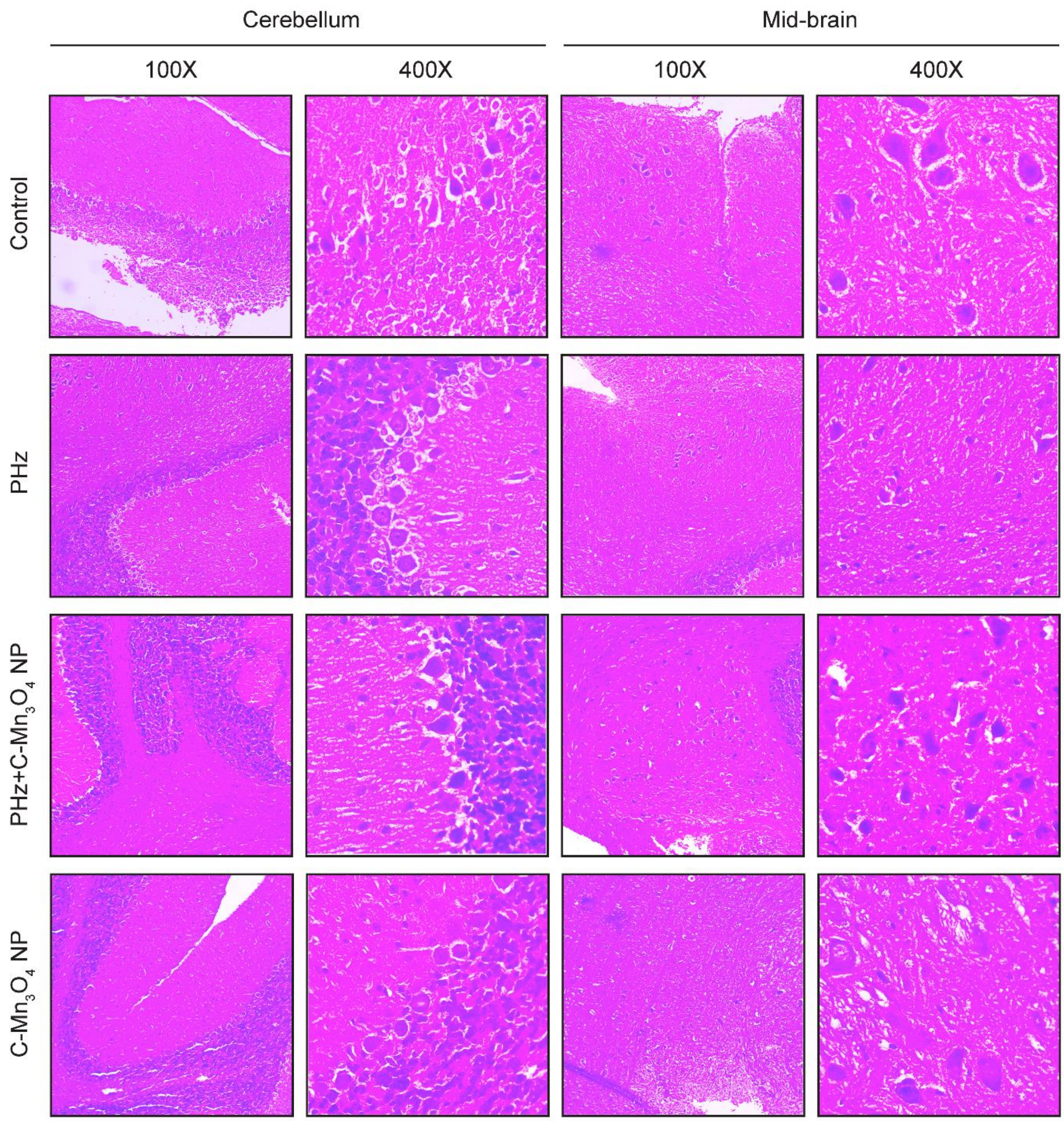


**Supplementary Figure S3. Micrographs of hematoxylin and eosin stained brain sections.**

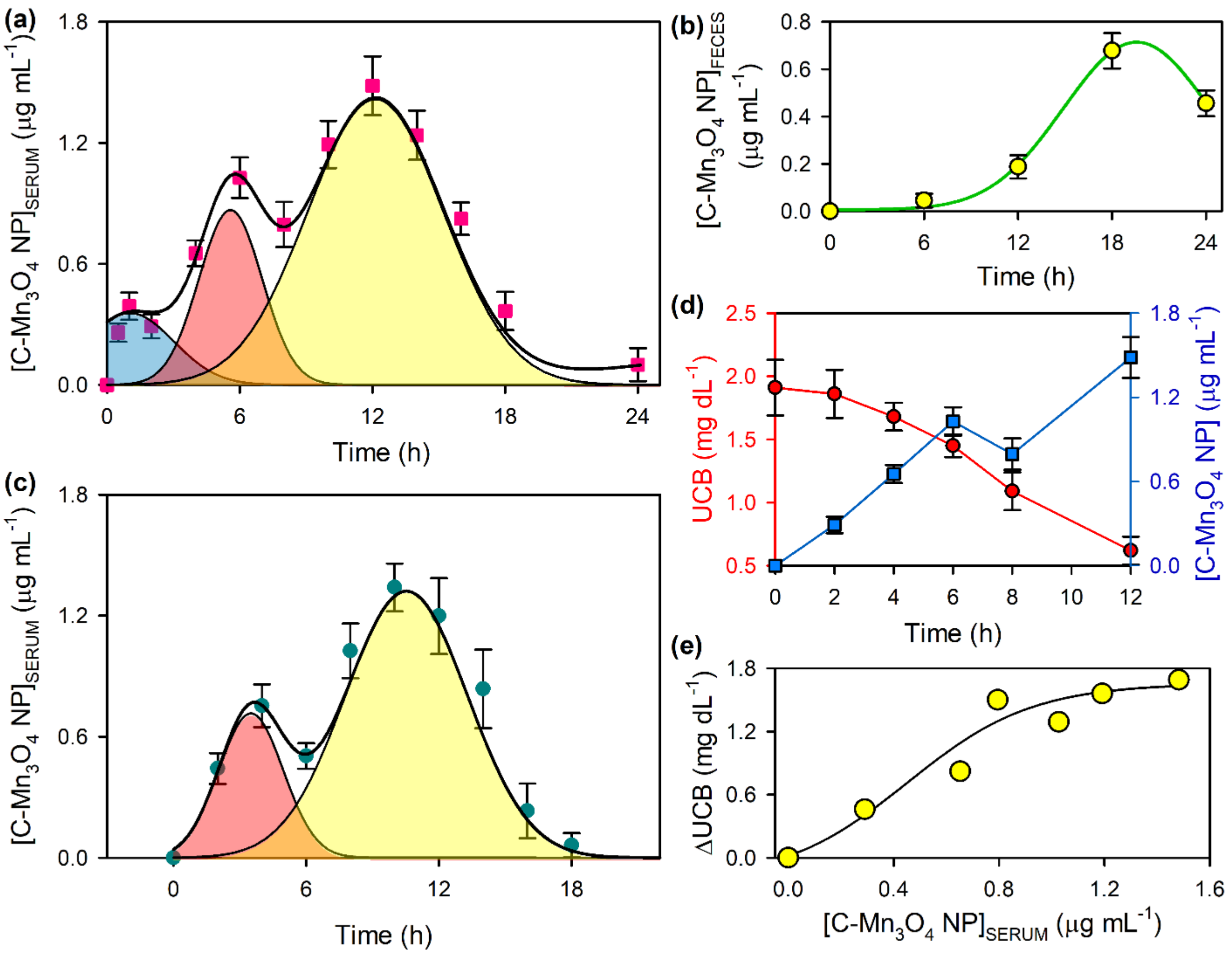


**Supplementary Figure S4. Pharmacokinetics and pharmacodynamics of C-$Mn_3O_4$ NPs.** (a) Plasma concentration-time profile following oral administration of NPs as measured using inductively coupled plasma-atomic emission spectroscopy (ICP-AES). The three peaks can be assigned to upper gastro intestinal (GI) tract absorption (peak 1), intestinal absorption (peak 2) and intestinal reabsorption (peak 3). (b) The elimination profile of C-$Mn_3O_4$ NPs through faeces (the main excretion route). (c) The first peak disappears in plasma concentration-time profile when administered directly to stomach. This confirms the upper GI tract absorption of NPs as the source of first peak. (d & e) Pharmacodynamics of bilirubin degradation.

**Supplementary Table S1: Pharmacological parameters of C-$Mn_3O_4$ NPs**

| **Parameters** | **Oral Gavage** | **Delivery to Stomach** |
|---|---|---|
| $C_{MAX}$ (mg $mL^{-1}$) | 1.71±0.25 | 1.84±0.21 |
| $t_{MAX}$ (h) | 12.1±0.2 | 10.4±0.2 |
| AUC (µg h $mL^{-1}$) | 18.01±1.47 | 16.08±2.17 |
| Clearance (L $h^{-1}$ $kg^{-1}$) | 12.3 | 11.3 |
| Bioavailability (%) | 12.2 | 11.9 |

All data represented as Mean ± Standard Deviation (SD). N=6 for each measurement.
